# A double-staining automated flow cytometry method for real-time monitoring of bacteria in continuous bioreactors

**DOI:** 10.1101/2025.03.24.644962

**Authors:** Juan López-Gálvez, Erik Schönfelder, Hanna Mayer, Konstanze Schiessl, Marisa O. D. Silva, Hauke Harms, Susann Müller

## Abstract

In industrial biotechnology, cell density is a critical parameter that influences key process control variables such as feed rate, harvest timing, and product recovery. Traditional biomass measurements are indirect, incapable of quantifying population heterogeneity, and often prone to inaccuracies due to manual handling and interference from culture media compounds. This study introduces an automated flow cytometry approach to enable continuous, real-time monitoring of bacterial cultures in a continuous bioreactor. Our method employs a double-staining protocol that combines DAPI for assessing total DNA content with Alexa Fluor 488 via Click-iT technology to detect the percentage of cells with active DNA replication through EdU incorporation. This streamlined protocol, which integrates cell fixation, permeabilization, staining, and measurement, was applied to three Gram-negative strains: *Bradyrhizobium sp.*, *Escherichia coli*, and *Stenotrophomonas rhizophila*. The approach successfully captured both growth dynamics and cell cycle progression, providing rapid and quantitative insights into culture composition. The enhanced temporal resolution achieved by this method facilitates timely adjustments of process parameters, ultimately improving bioprocess efficiency and product quality. These results underscore the potential of automated flow cytometry as a powerful tool for real-time monitoring and control in industrial biotechnology.

## 1. Introduction

Cell density is a fundamental parameter in industrial biotechnology, as it directly influences key process control variables such as feed rate, harvest time, and product recovery[1]. Monitoring and adjusting cell density are crucial for optimizing bioprocesses and achieving desired production outcomes[2]. Traditionally, biomass has been measured using bulk parameter analysis methods like dry or wet weight. These methods involve manual sampling and handling, making them imprecise, time-consuming, and susceptible to human error. Another common method is the measurement of optical density (OD), which estimates biomass by measuring light scattering; however, OD is only an indirect measure of cell concentration, and its accuracy can be compromised by interfering compounds in the culture media[3]. In addition, none of these tools is capable of quantifying population heterogeneity, a problem that can arise in bioprocesses, reducing overall bioprocess productivity[4]

As a result, relying solely on traditional biomass data and OD measurements can delay the acquisition of accurate information, making it challenging to adjust the process in real time and negatively impacting overall process efficiency. This situation emphasizes the need for more reliable and efficient monitoring methods. Advanced sensor technologies offer continuous, real-time data by directly measuring e.g., the dielectric properties of viable cells[5] or spectroscopic characteristics[6,7], thereby enabling improved process control in industrial biotechnology.

Automated flow cytometry has emerged as a powerful tool for microbial monitoring across diverse applications. This technique offers rapid, quantitative insights into bioreactor dynamics that are essential for optimizing bioprocesses[8,9]. Conventional flow cytometry is widely employed in biotechnology to measure variations in cell concentration and to distinguish different cell types within a productive microbial process[10]. Moreover, conventional flow cytometry has been extended to electricity generation processes, such as monitoring microbial communities in microbial fuel cells, where understanding community structure is vital for process optimization[11]. Additionally, researchers have applied flow cytometric techniques to study the microbiomes of rodents, providing insights into host– microbe interactions[12]. Beyond conventional biotechnology, automated flow cytometry is increasingly used for broader applications. It plays a critical role in drinking water microbiome analysis, where it helps in assessing water quality and microbial safety[13–15]. Automated flow cytometry has also been used effectively in the characterization of marine prokaryote communities in open water systems[16] .

In previous studies we have demonstrated the OC-300’s capability to establish automated flow cytometry procedures in conjunction with the CytoFLEX S. This combination has proven effective in delivering stable cell counts and facilitating the monitoring of bacterial cultures at temporal resolutions comparable to their generation times. Such high-resolution monitoring enables real-time insights into microbial dynamics, allowing for prompt detection of shifts in culture composition [8].

Cells exhibit a range of physiological states within a bioreactor that are largely governed by their cell cycle phases, and these differences can have a profound impact on cellular productivity[17,18]. In our study, we extend our analysis beyond quantifying changes in cell number by also investigating cell cycle dynamics. To this end, we employ two different dyes that allow us to differentiate between the various cell cycle phases by simultaneously assessing cell replication activity and cell cycle progression. The protocol eliminates centrifugation steps—including fixation, permeabilization, staining, and dilution—to facilitate on-line monitoring of growth in biotechnological systems.

DAPI (4’,6-diamidino-2-phenylindole) and Alexa Fluor 488 were used to simultaneously assess total DNA content and DNA synthesis. DAPI is a total DNA stain enabling for counting chromosome numbers of cells[19], whereas the Alexa 488 staining utilizes Click-iT technology, which employs EdU (5-ethynyl-2’-deoxyuridine), a thymidine analog present in the culture medium. EdU is incorporated into newly synthesized DNA strands during active DNA replication[20]. After incorporation, the EdU residue undergoes a copper-catalyzed click chemistry reaction, covalently binding to an Alexa Fluor 488 dye, which can then be detected by flow cytometry[21]. The Alexa 488 Click-iT technology is best known for the analysis of human and mammalian cultures but its usage in bacterial biotechnological applications has not been reported yet [22,23].

The study utilized three distinct Gram-negative strains: *Bradyrhizobium sp.*, *Escherichia coli*, and *Stenotrophomonas rhizophila*. By applying the double-staining technique to these strains within a continuous bioreactor, we explored whether cell growth and heterogeneous DNA replication dynamics could be effectively monitored and analyzed using on-line flow cytometry.

## 2. Material and Methods

### 2.1 Cultivation of bacterial cells

The strains *Bradyrhizobium* sp. Leaf396-mScar, obtained from Schlechter et al.[24], *Escherichia coli* K12 LE392 DSM 4230, and *Stenotrophomonas rhizophila* DSM 14405 (both obtained from the German Collection of Microorganisms and Cell Cultures (DSMZ), Leibniz Institute, Braunschweig, Germany) were initially cultivated on LB agar plates (Lysogeny Broth) at 30 °C for 72 h, starting from glycerol stock suspensions. Subsequently, 20 mL of liquid LB medium was inoculated with a single colony of each strain and incubated at 30 °C with shaking at 250 rpm for 24 h. After incubation, the optical density (OD_700 nm = 0.5 cm_) of the preculture was measured using an Ultrospec 1100 Pro spectrophotometer (Amersham Biosciences, Amersham, UK). The required volume to inoculate the main culture, whether for batch cultivation in a 24-well plate or for continuous culture in a bioreactor, was then calculated to achieve an initial OD_700 nm = 0.5 cm_ = 0.05.

### 2.2 Cultivation of bacterial cells in batch experiments

LB medium was used for these experiments. For experiments where Alexa 488 fluorescence was to be measured, 5-ethynyl-2′-deoxyuridine (EdU) was added to LB medium at an initial concentration of 3.75 or 7.5 µg / mL. The LB medium was then inoculated with sufficient preculture to achieve an initial OD_700 nm = 0.5 cm_ = 0.05. The culture was transferred to a 24-well plate, with each well containing 1 mL of the inoculated medium, and incubated at 30 °C with shaking at 150 rpm for 24 h for both *E. coli* and *Bradyrhizobium* sp., and 72 h for *S. rhizophila*. To ensure sterility while allowing air exchange, the plate was covered with a “Breathe-Easy” anti-evaporation foil (Merck KGaA, Darmstadt, Germany) placed in an Incubator Hood TH 30 (Edmund Bühler GmbH, Bodelshausen, Germany). Growth was measured by initial OD_700 nm = 0.5 cm_ = 0.05 hourly up to 8 h, and then at the 24 h mark and finally at 48 h and 72 h.

### 2.3 Cultivation of bacterial cells in a continuous bioreactor

EdU-LB medium was prepared at initial concentrations of either 3.75 or 7.5 µg/mL, and 10 mL of this medium was inoculated with an appropriate volume of preculture to reach an OD_700 nm=0.5 cm_ = 0.05. The bioreactor was maintained with a constant working volume of 10 mL at 30 °C and 250 rpm (Cimarec Poly 15 und Multipoint Stirrer, Thermo Fisher Scientific Inc., Waltham, MA, USA; Incubator Hood TH 120 25, Edmund Bühler GmbH, Bodelshausen, Germany). The inflow rate was set to 2.5 mL/h, the outflow rate to 1.3 mL/h (LabN1-II peristaltic pump, Drifton A/S, Denmark), and the sampling rate to 1.2 mL/h, with hourly sample collection. This configuration resulted in a dilution rate of 0.25 h⁻¹, corresponding to a total volume exchange time of 4 h. After 5 volume exchanges (20 h), the balance phase was considered to have been reached. For the D = 0.5 h^−1^ experiments, the inflow rate was set to 5 mL/h, the outflow to 3.8 mL/h and the sampling rate was kept the same. For the D = 0.31 h^−1^ experiment, the inflow and outflow rates were 3.125 mL/h and 1.925 mL/h, respectively. Oxygen concentration was monitored using the O_2_ Sensorspot SP-PSt6-YAU (PreSens Precision Sensing GmbH, Regensburg, Germany). Aeration was maintained by keeping a constant oxygen pressure at 40 psi, and a 0.2 µm filter (Labsolute, Renningen, Germany) was used for air exchange. The peristaltic tubing used was Tygon LMT-55, with a diameter of 1.295 mm (Saint-Gobain S.A., La Defense, France).

### 2.4 Paraformaldehyde (PFA)/EtOH fixation and DAPI staining for the DAPI fingerprint analysis

To perform a fingerprinting analysis of different subpopulations with varying chromosome numbers of pure strains, DAPI staining was carried out following PFA/EtOH fixation, as described by Cichocki et al.[25]. In short, cells from the batch cultures were sampled hourly and adjusted to an OD_700 nm = 0.5 cm =_ 0.5 in phosphate buffer (289 mM Na_2_HPO_4_ and 128 mM NaH_2_PO_4_ with double-distilled water, pH 7), then centrifuged at 3200× g at 4°C for 10 min. The cells were incubated for 30 min in 2 mL of a 2% PFA solution at room temperature (RT). After incubation, the PFA was removed by centrifugation (3200× g, 4°C, 10 min), and the cells were resuspended in 2 mL of a 70% EtOH fixation solution for at least 1 h at RT. The fixed cells were then washed with PBS buffer by centrifugation (3200× g, 4°C, 10 minutes) and adjusted to 2 mL with an OD_700 nm = 0.5 cm_ = 0.04. The PBS was discarded by centrifugation (3200× g, 4°C, 10 min), and 2 mL of a 0.24 µM DAPI staining solution in PBS buffer, diluted from a DAPI stock solution (143 µM DAPI dissolved in 100 µl dimethylformamide and then in double-distilled water) was added and incubated overnight. After incubation, the sample was ready to be measured.

### 2.5 DAPI and Alexa-488 double staining procedure for cell concentration determination and DNA synthesis activity

DAPI and Alexa-488 (Thermo Fisher Scientific Inc., Waltham, MA, USA) staining were used to assess cell number and DNA synthesis activity. This process required fixation and permeabilization to ensure effective staining. The fixation method employed was a previously validated NaCl/NaN_3_ /EtOH protocol[8].

The NaCl/NaN_3_/EtOH fixation was carried out by adding 475 µL of 30% NaCl, 35 µL of 20% NaN_3_ (both Merck KGaA, Darmstadt, Germany), and 100 µL of 70% EtOH (Chemsolute, Renningen, Germany) to 100 µL of the bacterial sample. This resulted in final concentrations of 20% NaCl, 1% NaN_3_, and 10% EtOH, with an incubation time of 10 min. Following fixation, a permeabilization step using Triton X (Merck KGaA, Darmstadt, Germany) was carried out. Specifically, 200 µL of 0.5% Triton X was added to 100 µL of the previously fixed cell suspension and incubated for 20 min.

Subsequently, the double staining with DAPI and Alexa-488 was carried out simultaneously. This entire procedure was either performed manually by pipetting or automatically by the OC-300 automation unit (onCyt Microbiology, Zürich, Switzerland) (Figure 1). DAPI staining was achieved using a 1 µM DAPI staining solution in PBS buffer. The Alexa-488 stain was performed using the “Click-iT EdU Cell Proliferation Kit for Imaging, Alexa Fluor™ 488 dye” (catalogue number C10337 Thermo Fisher Scientific Inc., Waltham, MA, USA). From the kit the following amounts of chemicals were used to prepare one volume of reaction cocktail:

**Figure 1.**
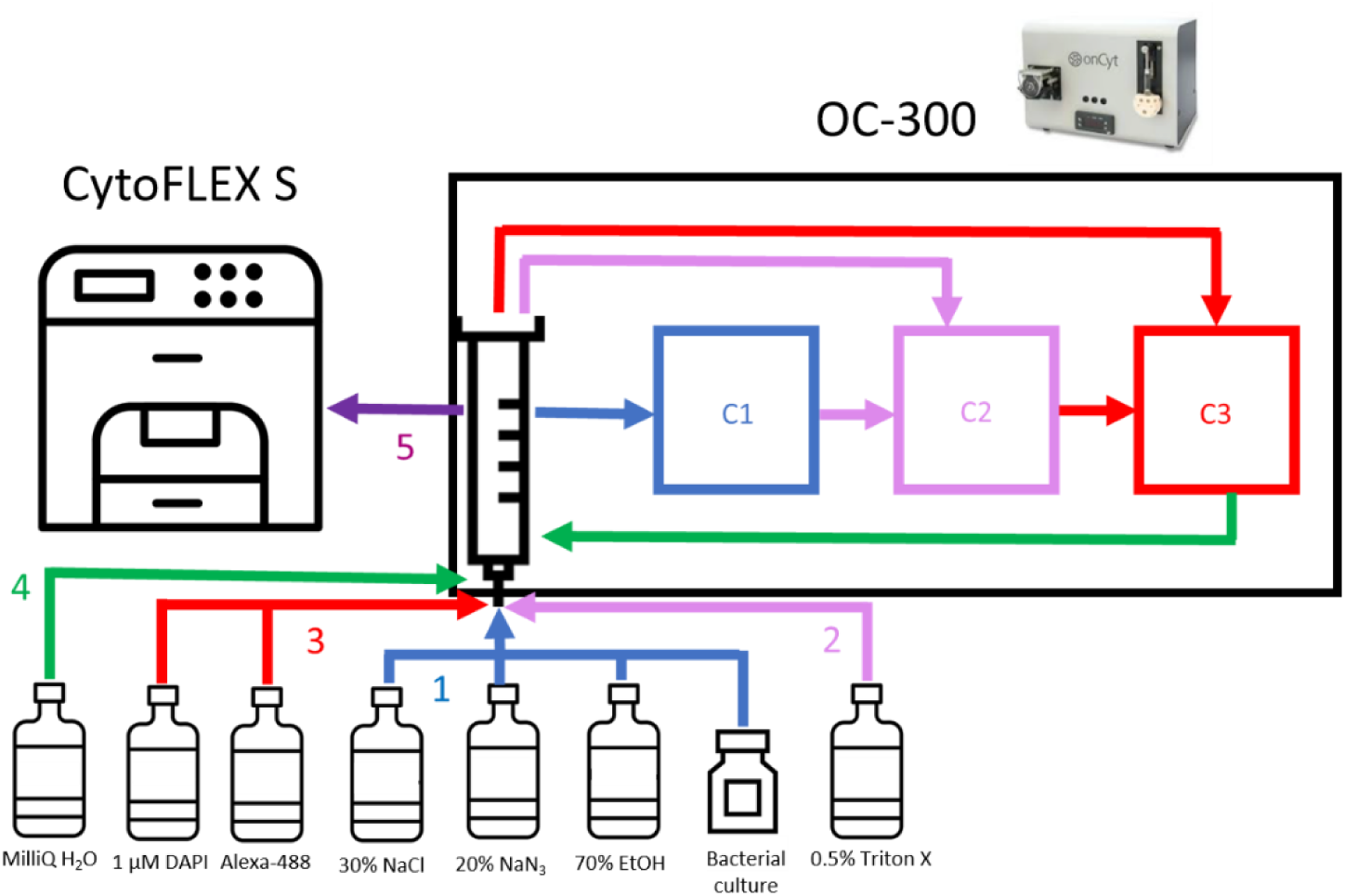
A schematic representing the workflow of automated fixation, permeabilization, double staining, dilution and measurement performed by the OC-300. (1) The sample drawn from the bacterial culture was then moved to chamber 1 (C1) and fixed with NaCl (20%), NaN_3_ (1%), and EtOH (10%) for 10 min. (2) The fixed sample was moved from C1 to chamber 2 (C2) and permeabilized with 0.5% Triton X for 20 min. (3) After permeabilization, the sample was transferred from C2 to chamber 3 (C3) and stained with DAPI (1 µM) and Alexa-488 reaction cocktail for 10 min. (4) The stained sample was diluted 1:20 with MilliQ water in the syringe. (5) Finally, the diluted sample was sent to the cytometer for measurement.

Once the Alexa-488 staining solution had been prepared, the double staining procedure was performed. To do this, 200 µL of 1 µM DAPI staining solution and 40 µL of the Alexa-488 reaction cocktail were added to 100 µL of the permeabilized cell suspension and incubated for 10 min. After the incubation, the stained cell suspension was diluted 1:20 in MilliQ water (IQ 7000 Ultrapure Lab Water System, Merck KGaA, Darmstadt, Germany) and measured immediately.

### 2.6 Automated workflow for the on-line double staining of a bacterial culture

The double staining procedure outlined above can be automated and performed on-line by the OC-300 automation unit. In this process, the OC-300 drew a sample from either the batch culture or the continuous bioreactor, and then carried out the sequential steps of fixation, permeabilization, double staining, and final dilution. Once these steps were completed, the stained cell suspension was sent to the CytoFLEX S flow cytometer (Beckman Coulter, Brea, CA, USA) for measurement. After each measurement, the device cleaned itself and prepared for the next sample. Considering the fixation, permeabilization and staining times, and the cleaning process between samples, a new sample was drawn and measured every 60 min.

### 2.7 OC-300 automation unit

The OC-300 automation unit was engineered for the automated flow cytometry analysis of bacterial communities in both technical and environmental water systems, including fresh water, drinking water, and wastewater [14,15]. It features two valves, each with twelve ports, that allow for the intake and dilution of samples, as well as the addition of reagents for fixation, permeabilization, and staining via a syringe. These valves also link the incubation chambers within the device. Inside the three incubation chambers, bacterial samples can be mixed with various solutions to carry out fixation, permeabilization, or staining steps. The unit was controlled by the cyOn software (onCyt Microbiology, Zürich, Switzerland) and was connected to the bioreactor through a sampling port, while the flow cytometer was linked via the OC-300-CytoFLEX interface. In the case of 24-well plate experiments, the sampling tubing was inserted into the well at every sampling time.

### 2.8 CytoFLEX S

A CytoFLEX S flow cytometer (Beckman Coulter, Brea, CA, USA), operated with the CytExpert software (Beckman Coulter, Brea, CA, USA), was employed for this study. The instrument is equipped with 375 nm (60 mW), 488 nm (50 mW), and 638 nm (50 mW) lasers. The 488 nm laser was used to detect forward scatter (FSC) (488/8 nm band-pass), side scatter (SSC) (488/8 nm band-pass, trigger signal), and Alexa-488 fluorescence (525/40 nm band-pass). The DAPI fluorescence (450/45 nm band-pass) was measured using the 375 nm laser for excitation. The fluidic system was operated at a constant speed of 60 µL/min. For optical calibration in the logarithmic range, 0.5 µm and 1.0 µm UV Fluoresbrite microspheres (Polysciences, Cat. Nos. 18339 and 17458, Warrington, PA, USA) and 0.5 µm and 1.0 µm Yellow Green Fluoresbrite microspheres (Polysciences, Cat. Nos. 17152-10 and 17154-10, Warrington, PA, USA) were used.

### 2.9 Bioinformatic tools

The cell concentration, relative subpopulation proportions and relative DNA synthesis activities were calculated using the software FlowJo (BD Biosciences, Franklin Lakes, NJ, USA). In this software, gates were defined to include cells with similar characteristics that group together as a subpopulation. FlowJo was also used to create the flow cytometric 2D plots. The creation of the barcode images for the analysis of different chromosome number subpopulations was done by the flowCyBar software[11] embedded in the biTCa Analyze Tool graphical user interface (GUI) developed by Bruckmann et al. (2022) [26]. The time series graphs were made in the software Origin 2023 (OriginLab Corporation, Northampton, MA, USA).

## 3. Results

The most important parameter to obtain information about bacterial activity is the measurement of growth. In a batch culture this is usually performed by OD measurements, which were also carried out in this study. However, OD analysis has several disadvantages: It is a bulk parameter (no single-cell information captured), it also measures colors and particles of the medium and ignores changes in bacterial cell sizes. In contrast, on-line flow cytometry tracks growth at the individual cell level and provides exact cell numbers per volume. In addition, in this study, a DNA fluorescent dye was used to obtain information about proliferation states of bacteria. The best-known dye for this purpose is DAPI, which binds AT-specifically to the DNA of bacteria. The amount of DNA per cell was reflected by the height of the fluorescence intensity (DAPI-FI) at the blue range of the light after excitation of the DAPI stained cells with a 375 nm laser. An automated cytometric on-line approach based on DAPI-labelled single cell analysis is available and has been previously validated for analytical reliability[8]. This technology has now been coupled with a bioreactor system to document on-line bacterial growth under balanced continuous cultivation conditions.

### 3.1 On-line cytometry as a sensor for bacterial growth

Three different strains were selected to be grown under continuous balanced growth conditions. The strains were pre-cultivated separately as described in the method section and bioreactors were inoculated to an OD_700nm =0.5 cm_ = 0.05, respectively. The continuous bioreactor system consisted of a 10 mL bioreactor connected both to the medium source and the automatic sampler OC-300 and was run with a dilution rate D=0.25 h^−1^ and kept constant at T= 30°C, at 250 rpm and an oxygenation pressure of 40 psi. The bioreactor was sampled and measured automatically every hour and oxygen concentration was measured in parallel. A 5-fold volume exchange was reached after 20 h to guarantee balanced conditions in the bioreactor system.

During adaptation to the bioreactor conditions, the strain *Bradyrhizobium* sp. immediately started to grow which continued until 18 h when the culture transitioned into the balanced phase, stabilizing at 4.35 × 10^9^ ± 2.12 × 10^8^ cells/mL (Figure 2A). The period of exponential growth was accompanied by DNA synthesis, as shown by the DAPI mean fluorescence intensity (FI) curve reaching maximal FI values of around 3.5 × 10^4^ [rel. units] DAPI-FI at 6 h. After the DAPI-FI gradually decreased until it balanced at 1.94 × 10^4^ ± 1.74 ×10^3^ [rel. units] at 18 h. The high growth activities were accompanied by a decrease in oxygen concentration to 1.65 ± 0.28 mg/mL in the balanced phase.

**Figure 2.**
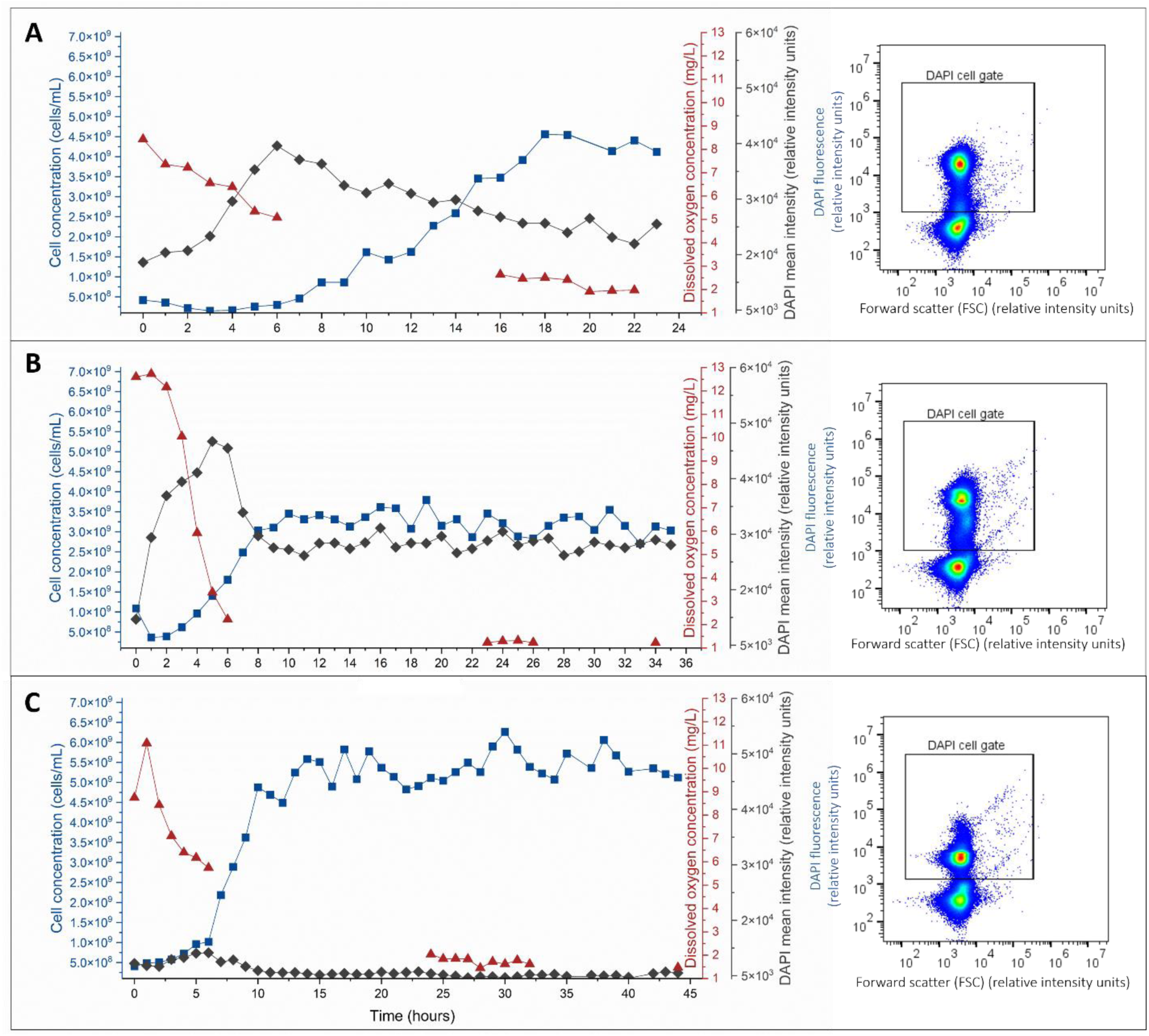
Automatic on-line cytometric analysis of cell growth in 10-mL bioreactors of 3 different strains under continuous balanced conditions. Cell concentration (dark blue), dissolved oxygen concentration (red) (measured manually), and DAPI mean fluorescence intensity (dark gray) was hourly measured at conditions of D= 0.25 h^−1^, T=30°C, and 250 rpm. **A:** *Bradyrhizobium* sp. **B:** *E. coli* and **C:** *S. rhizophila*. The right 2D plots show examples of DAPI stained cells vs. forward scatter as an example at 20 h cultivation.

The strain *E. coli* showed a different behavior (Figure 2B). Even though this bioreactor was also inoculated to the same OD of 0.05, growth started faster and balanced already after 8 h at around 3.24 × 10⁹ ± 2.63 × 10^8^ cells/mL. The high proliferation and uncoupled DNA synthesis during the first hours is visualized by the massive increase in DAPI-FI to 4 × 10^4^ [rel. units] during the adaptation phase until it levels to clearly lower values in the balanced phase (2.34 × 10^4^ ± 1.74 × 10^3^ [rel. units]). The period of adaptation to the bioreactor conditions was accompanied by a manifest decrease in dissolved oxygen concentration to nearly zero, reflecting the high metabolic activity of the cells during rapid proliferation.

The strain *S. rhizophila* shows a similar fast adaptation to reactor conditions (Figure 2C). Starting also with an of OD_700nm =0.5 cm_ = 0.05, adaptation to reactor conditions was rapid and quickly transitioned into exponential growth until it reached the balanced phase already at 10 h, at a cell concentration of 5.34 × 10⁹ ± 4.06 × 10^8^ cells/mL. The almost double cell number compared to the *E. coli* strain was expected because the single cell size of *S. rhizophila* is much smaller in comparison and therefore the bioreactor was inoculated with a higher initial cell number. The available nutrients were apparently sufficient to produce higher cell numbers compared to *E. coli* as was already described by Cichocki et al.[25]. The DAPI-FI values were significantly lower than those of the other strains at about 5.19 × 10^3^ ± 3.10 × 10^2^ [rel. units] due to the lower DNA content per cell, but peaked similarly during the adaptation phase. As with the other strains, a decrease in dissolved oxygen levels was observed, corresponding to the increased metabolic activity during cell growth.

In order to prove the sensitivity of the on-line cytometric analysis procedure, the three strains were additionally all grown at a higher dilution rate of D = 0.5 h^−1^. This rate was higher than the estimated µ_max_ values for each strain, which were determined to be 0.28 h^−1^ for *Bradyrhizobium* sp., 0.30 h^−1^ for *E. coli*, and 0.31 h^−1^ for *S. rhizophila* in batch cultivations of the same medium. Therefore, a washout of the 3 strains was expected. Instead, cells were still detected (although at a lower concentration) in the balanced phase accompanied by still high DAPI-FI values (Table 2).

**Table 1.** Click-iT reaction cocktail of Alexa Fluor-488 staining solution. Volumes for 1 sample.

| Reagent | Volume (μL) |
| --- | --- |
| 1x Click-iT EdU reaction buffer | 34.4 |
| Copper sulfate (100 mM) | 1.6 |
| Alexa Fluor 488 picolyl azide (10 mM) | 0.096 |
| Click-iT EdU buffer additive | 4 |

**Table 2.** Summary of the cell concentration (cells/mL), DAPI fluorescence intensity (relative intensity), and oxygen concentration (mg/L) recorded during the balanced phase at different dilution rates for the three strains examined in Figures 2, 3, and SI 1.

|  | dilution rate (h<sup>-1</sup>) | cell concentration (cells / mL) | DAPI-FI [rel. units] | oxygen concentration (mg / L) |
| --- | --- | --- | --- | --- |
| *Bradyrhizobium* sp. | 0.25 | $4.35 \times 10^9 \pm 2.12 \times 10^8$ | $1.94 \times 10^4 \pm 1.74 \times 10^3$ | $1.65 \pm 0.28$ |
| | 0.5 | $1.08 \times 10^9 \pm 1.51 \times 10^8$ | $2.50 \times 10^4 \pm 3.44 \times 10^3$ | $1.57 \pm 0.02$ |
| *E. coli* | 0.25 | $3.24 \times 10^9 \pm 2.63 \times 10^8$ | $2.34 \times 10^4 \pm 1.17 \times 10^3$ | $0.76 \pm 0.04$ |
| | 0.31 | $2.70 \times 10^9 \pm 3.03 \times 10^8$ | $2.90 \times 10^4 \pm 2.00 \times 10^3$ | $0.53 \pm 0.03$ |
| | 0.5 | $1.19 \times 10^9 \pm 1.93 \times 10^8$ | $3.96 \times 10^4 \pm 6.38 \times 10^3$ | $0.21 \pm 0.01$ |
| *S. rhizophila* | 0.25 | $5.34 \times 10^9 \pm 4.06 \times 10^8$ | $5.19 \times 10^3 \pm 3.10 \times 10^2$ | $1.23 \pm 0.18$ |
| | 0.5 | $1.56 \times 10^9 \pm 1.41 \times 10^8$ | $1.43 \times 10^4 \pm 1.69 \times 10^3$ | $1.56 \pm 0.34$ |

In the case of *Bradyrhizobium* sp. (Figure 3A), the adaptive growth began 4 h after the start of the experiment and lasted 8 h, reaching the equilibrium phase earlier compared to D = 0.25 h^−1^, but maintaining only a cell concentration of 1.08 × 10⁹ ± 1.51 × 10^8^ cells/mL. The DAPI-FI values increased also faster, peaked at 4h with DAPI-FI of about 3.3 × 10^4^ [rel. units] and reached a level of 2.5 × 10^4^ ± 3.44 × 10^3^ [rel. units] in the balanced phase (Table 2). The increase in DAPI-FI was accompanied by a decrease in oxygen concentration.

**Figure 3.**
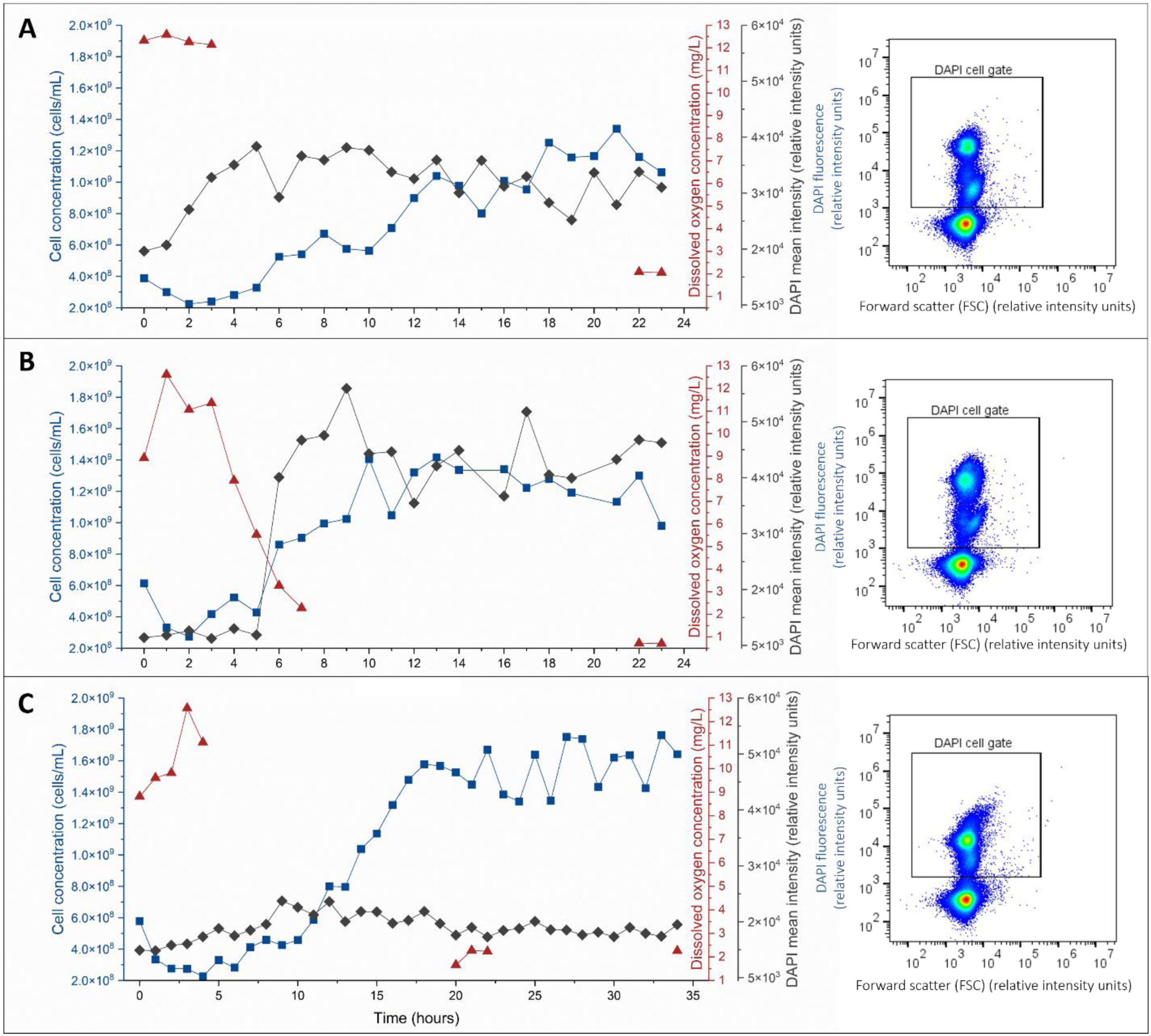
Automatic on-line cytometric analysis of cell growth in 10-mL bioreactors of 3 different strains under continuous balanced conditions for the determination of the replication activity. Cell concentration (dark blue), dissolved oxygen concentration (red) (measured manually) and DAPI mean fluorescence intensity (dark gray) were measured hourly at conditions of D= 0.5 h^−1^, T=30°C, and 250 rpm. **A**: *Bradyrhizobium* sp. **B**: *E. coli* and **C**: *S. rhizophila*. The 2D plots to the right show examples of DAPI stained cells vs. forward scatter as an example at 20 h cultivation.

For *E. coli* (Figure 3B), a similar pattern was observed, with a 4-h lag phase before the onset of adaptive growth, which lasted for 6 h. At the following balanced conditions, a cell concentration of approximately 1.19 × 10⁹ ± 1.93 × 10^8^ cells/mL was reached which was only a third of the value obtained for the D = 0.25 h^−1^. The DAPI-FI values peaked at 9 h and was also clearly higher compared to the dilution rate before with approximately 4.1 × 10 ^4^ [rel. units]. The DAPI-FI values stabilized at the double amount of around 3.96 × 10^4^ ± 6.38 × 10^3^ [rel. units] compared to the lower dilution rate. The oxygen values reached a concentration near zero. The same experiment was also conducted at an intermediate dilution rate of D = 0.31 h^−1^ which is near the µ_max_ value. Also, under these conditions the cell concentrations were similar to those of D = 0.25 h^−1^ with only a slight decrease to 2.70 × 10^9^ ± 3.03 × 10^8^ cells/mL. The same is valid for the DAPI-FI values with 2.90 × 10^4^ ± 2.00 × 10^3^ [rel. units] (Table 2, SI Figure 1). In addition, the oxygen consumption was in the same range.

For *S. rhizophila* (Figure 3C), a longer lag phase of 10 h was observed before the onset of exponential growth phase which lasted 8 h. Under balanced conditions, only one-third of the cell concentration achieved under D = 0.25 h^−1^ was reached, with a value of 1.56 × 10⁹ ± 1.41 × 10^8^ cells/mL. The DAPI-FI showed a continuous increase from the start of the experiment until the peak was reached at 9 h during the onset of the adaptive phase with 1.9 × 10^4^ [rel. units]. In the balanced phase the mean DAPI-FI was lower with 1.43 × 10^4^ ± 1.69 × 10^3^ [rel. units] but reached a 5-fold value compared to the conditions of D = 0.25 h^−1^ (Table 2). The provided oxygen was not fully used and analyzed to be 1.56 ± 0.34 mg/mL dissolved oxygen concentration.

### 3.2 Determination of the replication activity percentage

Automated on-line cytometry also allows the use of many other fluorescent markers. For example, on-line cytometry has been used for many years to follow the dynamics in oceans and surface waters based on the autofluorescent properties of autotrophic microorganisms in these environments[16]. Furthermore, it can be used to automatically analyze the presence of heterotrophic organisms when stained with nucleic dyes[14,15]. In addition, the expression of intrinsic genetic markers can also be detected[9]. In this study, however, we were particularly interested in following bacterial growth in bioreactor systems. In addition to DAPI which labels chromosomes per cell, we have introduced an Alexa 488 dye that specifically labels DNA-replicating cells. Using the Click-iT technology, EdU is added to the medium and incorporated into the DNA when cells replicate. In a next step, an Alexa 488 azide derivative is added to the cell solution shortly before measurement to specifically label the EdU nucleic acid in the DNA. This step is facilitated by the use of copper as a catalyst.

The on-line cytometric procedure developed in this study is new and involves for the first time two fluorescent dyes (DAPI and Alexa 488). The procedure comprises the sampling from the bioreactor, fixation, DAPI staining, Alexa 488-staining, and precise dilutions steps using the OC-300 device. There are no centrifugation steps in between. The DAPI and Alexa 488 double staining of bacterial samples was used both to determine cell concentration and DNA replication percentage over time. The DNA-replication percentage was determined in the following way:

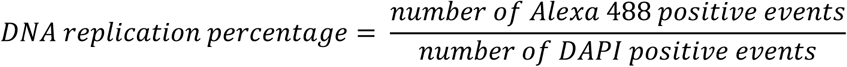

**Formula 1**. Calculation of DNA replication percentage. The number of total events in both the Alexa 488 and DAPI cell gates is determined and the former is divided by the later to obtain the DNA replication percentage.

To validate this method, we conducted in a first step a series of batch cultures, which were automatically sampled and measured. The cultures included individual strains of *Bradyrhizobium sp.*, *E. coli*, and *S. rhizophila*, respectively, grown in 24-well plates at 30°C and 150 rpm. Additionally, 100 µL samples were taken bihourly to measure OD at 700 nm. The results are shown in Figure 4.

**Figure 4.**
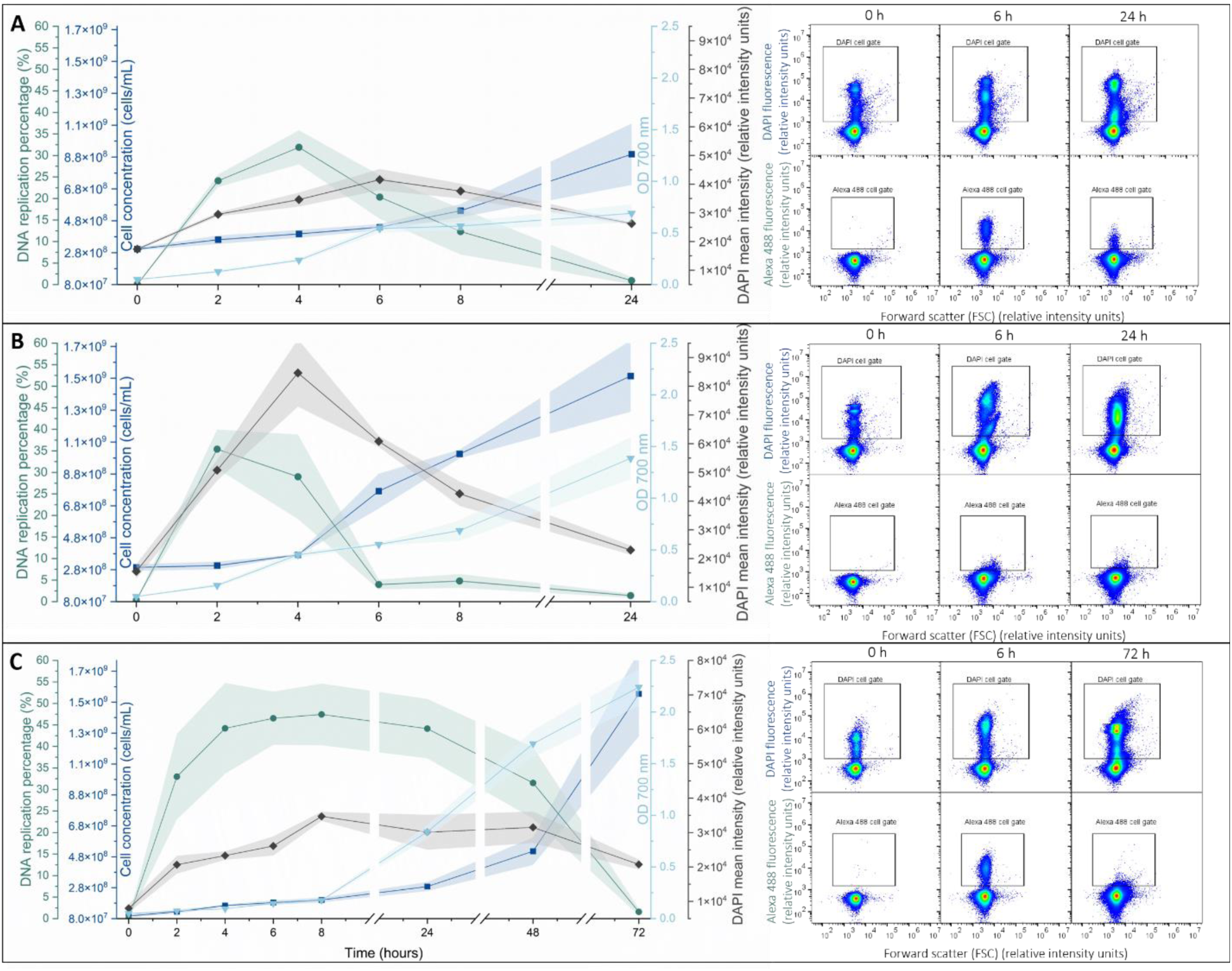
Automatic cytometric analysis of cell growth of 3 different strains. Cells were batch-cultivated in 24-well plates and bihourly sampled by the OC-300. OD (light blue), cell concentration (dark blue), DAPI replication percentage (green), and DAPI mean fluorescence intensity (dark gray) were measured bihourly. **A**: *Bradyrhizobium* sp. (addition of 7.5 µg/mL EdU) **B**: *E. coli* (addition of 3.75 µg/mL EdU) and **C**: *S. rhizophila* (addition of 3.75 µg/mL EdU). The right 2D plots show examples of DAPI stained cells (above) or Alexa 488 (below) vs. forward scatter at 0 h, 6 h, and 24 h respective 72 h cultivation. Points show average value and shaded area shows standard deviation.

Figure 4 illustrates that our automated procedure successfully performed the double staining procedure, allowing for determination of cell concentrations for all three strains. In addition, DAPI-FI values and DNA replication percentage values were easily recorded. In the case of *Bradyrhizobium* sp. And *E. coli* (Figure 3A,B), the percentage of DNA replication showed a sharp and immediate increase directly after inoculation, reaching the peaks at 4 h and 2 h, respectively. After these points, the replication activity gradually declined, eventually dropping to nearly undetectable levels at 24 h. This trend suggests an initial phase of rapid cellular replication activity followed by peak DAPI-FI postponed by an hour, respectively. A slowdown in growth activity was observed over the following hours as the bacterial cells entered the stationary phase. Consistent with these data, the cell concentrations showed a slow but steady increase throughout the observation period, starting at 3.03 × 10⁸ ± 8.41 × 10^6^ cells/mL at 0 h and rising to 8.97 × 10⁸ ± 1.21 × 10^8^ cells/mL at 24 h for *Bradyrhizobium* sp. and starting at 2.94 × 10⁸ ± 2.08 × 10^7^ and rising to 1.49 × 10^9^ ± 2.23 × 10^8^ cells/mL for *E. coli* (Figure 4AB).

In the case of *S. rhizophila* (Figure 4C), the DNA replication percentage increased also rapidly during the first 2 h and then remained high and unchanged throughout the following hours, persisting even up to 48 h. It is only after 72 h of culture that DNA replication activity dropped to nearly zero. Interestingly, once replication ceases at 72 hours, a sharp increase in cell concentration was observed. This suggests increased cell division activities at the end of the log-phase of growth, resulting in a sudden rise in cell number to 1.53 × 10^9^ ± 2.74 × 10^8^ cells/mL. This correlation between DNA replication and cell concentration highlights the effectiveness of the automated procedure in capturing dynamic bacterial growth patterns over time.

Having demonstrated that the double staining procedure effectively captured the behavior of the three different bacterial strains and tracked their changes in DNA replication percentage over time, the next step was to integrate this procedure into a continuous reactor system where the OC-300 and the CytoFLEX S were connected to the bioreactor and the automated sampling process was integrated.

We chose the D = 0.25 h^−1^ to follow growth below µ_max_ values to avoid the washout points and to guarantee balanced growth conditions. As before, cell number, DAPI-FI, and DNA replication percentage were determined automatically using the OC-300 connected to the CytoFLEX S and the controlled continuous bioreactor system. EdU was added to the culture media at a concentration of 3.75 µg/mL for all three strains. The oxygen concentration was measured as a standard. The automated procedure then sampled and performed the double staining of the bacterial cells every hour as before.

In the experiment with *Bradyrhizobium* sp. (Figure 5A), we observe that the cell concentration remained low but stable in the initial hours, without a significant increase. It slowed even down until 20 h, but finally entered a short growth phase of 5 h. During this period, the cell concentration recovered and reached the initial inoculation value of 6.31 × 10⁸ ± 1.07 × 10^7^ cells/mL. Oxygen consumption followed the same pattern. No oxygen was consumed during the decrease in cell number, but decreased slightly as growth increased after 20 h. The DAPI-FI values supported these data. However, the DNA replication percentage showed a different behavior. While it was relatively high during the first 7 h at about 15%, it decreased to about 5% and after 20 h to even 2%, where it stabilized for the rest of the experiment. This indicates a very low proliferation activity.

**Figure 5.**
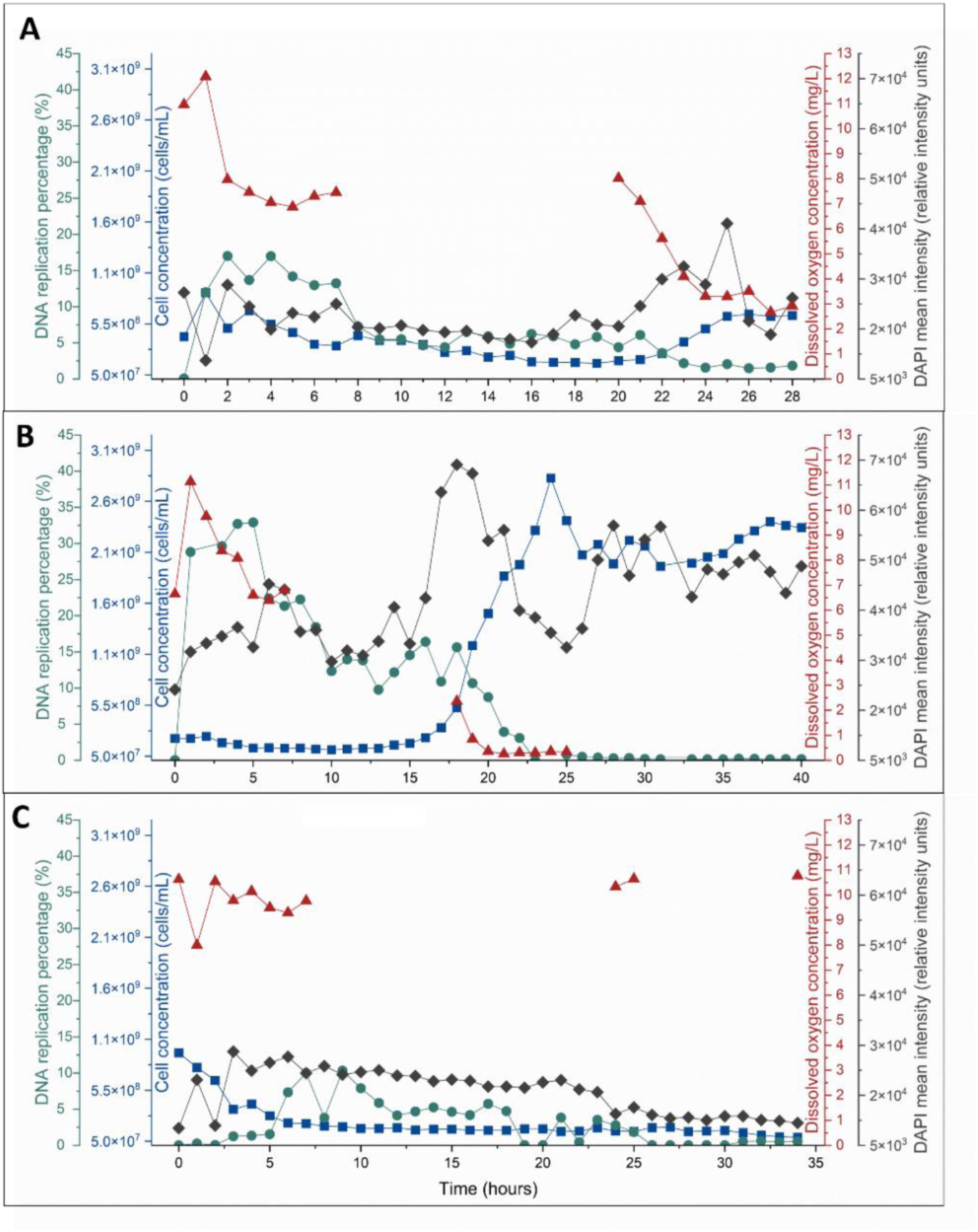
Automatic on-line cytometric analysis of cell growth in 10-mL bioreactors of 3 different strains under continuous balanced conditions for the determination of the replication activity. Cell concentration (dark blue), dissolved oxygen concentration (red) (measured manually), DAPI mean fluorescence intensity (dark gray) and DNA replication percentage (green) were measured hourly at conditions of D= 0.25 h^−1^, T=30°C, and 250 rpm. **A:** *Bradyrhizobium* sp. **B:** *E. coli* and **C:** *S. rhizophila*.

In the case of *E. coli* (Figure 5B), we observed a different behavior. The DNA replication percentage reached the highest value within the first 3 h of the experiment (33%), then decreased slowly but continuously over the following hours, dropping to around 5% after 20 h. During the first 20 h, no significant increase in cell concentration was observed although the DAPI-FI increased sharply starting at 15 h from 3.3 × 10^4^ [rel. units] to a peak value at 17 h with 6.4 × 10^4^ [rel. units]. Only at the 20-h mark, the massive exponential growth phase became apparent, lasting for 5 h, during which the cell concentration increased rapidly, reaching the balanced phase with 2.78 × 10⁹ ± 1.14 × 10^8^ cells/mL at 25 h. The balanced phase was accompanied by a significant decrease in DNA replication percentage, stabilizing only at around 0.5%. This behavior was similar to *Bradyrhizobium* sp. (Figure 5A), where a period of relatively high DNA replication was followed by a decrease. Different from the *Bradyrhizobium* sp. the oxygen concentration was nearly zero, pointing still to active growth.

In the case of *S. rhizophila* (Figure 5C), both cell concentration and DNA replication activity slowly reached the washout point. No oxygen was consumed. After a brief increase in cell number during the first 5 h, accompanied by a similar brief increase in DNA replication activity, all growth seemed to slow down. Also, in this bioreactor with *S. rhizophila,* both abiotic and cellular parameters were not in the equilibrium.

## 4. Discussion

We have shown that automated on-line flow cytometry allows the monitoring and control of bioreactor systems, offering real-time insights into cell populations and their dynamics. Unlike traditional offline sampling methods, which are time-consuming and can introduce variability and delay, on-line cytometry provides continuous data collection, enabling precise adjustments to optimize growth conditions. This technology enhances process efficiency by detecting changes in cell concentration, cell activity, size, and DNA content, allowing for immediate process control actions. As bioprocessing industries move toward more sophisticated, scalable and automated production methods, on-line cytometry can play a vital role in ensuring maximum production as well as consistent product quality and process stability[27].

Cell number and DAPI fluorescence intensity (DAPI-FI) were proven to serve as highly effective control parameters for monitoring cell growth in bioreactor systems, while bulk parameters such as optical density (OD) and dissolved oxygen concentration provide more general insight into overall culture conditions[28,29]. An early increase in DAPI-FI as was shown in Figures 2 and 3 for all three strains is a strong early indicator of cell proliferation. DAPI-FI was also the first signal to increase in the batch cultures (Figure 4). The rise in DAPI-FI can be seen as an early, sensitive signal of the onset of DNA replication and the beginning of cell cycle progression, followed by the readiness of the cells to divide. This early growth indicator precedes the increase in cell number, providing an opportunity for timely intervention and decision making. On the other hand, when cells have entered the balanced growth phase in a bioreactor, the proportion of cells actively replicating their DNA can only be approximated by the DAPI-FI values. Therefore, the use of a dye such as Alexa 488 to label the EdU incorporated into the DNA is a valuable complementary tool.

The Alexa 488 EdU staining procedure was originally developed to mark the replication of DNA in human cell lines[22,30]. To adapt the method for bacteria, we tested over ten Gram-positive and Gram-negative strains using the on-line staining procedure (List SI3), but with limited success: no Alexa 488 fluorescence could be detected. However, *E. coli* proved to be particularly amenable to the method, as we demonstrated effectively when cells were fixed using a standard protocol[25] that included centrifugation steps (SI Figure 2). In addition, the Alexa 488 signal was more readily detected than the DAPI-FI signal, also when automated on-line flow cytometry was used. Similarly, *Bradyrhizobium* sp. and *S. rhizophila* (Figure 4) appeared promising. However, while *E. coli* showed a significant response to the Alexa 488 EdU dye in the continuous bioreactor experiment (Figure 5B), the other two strains did not. But also *E. coli* showed a delayed growth response for both DAPI-FI and cell number and exhibited a long adaptation period (compared to the same experimental conditions without EdU, Figure 3). It appears that the presence of EdU significantly affected the growth of the bacteria in the continuous bioreactor. The growth of *S. rhizophila* (Figure 5C) was most severely inhibited, preventing sufficient growth to reach equilibrium, resulting in washout. It has been described that the incorporation of EdU inhibit the growth of both human and bacterial cells[31,32]. Independently of these results, we also tested whether a limitation of the Alexa 488 concentration underestimated the replication activities of the cells by growing *E. coli* at D= 0.25 h^−1^. The Alexa 488 dye was added up to the tenfold during the balanced phase of growth, but only 5% of the cells could be labelled. After a long adaptation period with almost no growth (and no oxygen consumption), *E. coli* reached almost the same DAPI-FI and cell number values as before (Figure 6). However, due to the limited applicability of the Click-it reaction to multiple strains and the high cost, it is recommended that Alexa 488 EdU not be used for routine DNA replication percentage determination. However, the procedure demonstrated that a double staining approach can be effectively managed using the OC-300 when necessary.

**Figure 6.**
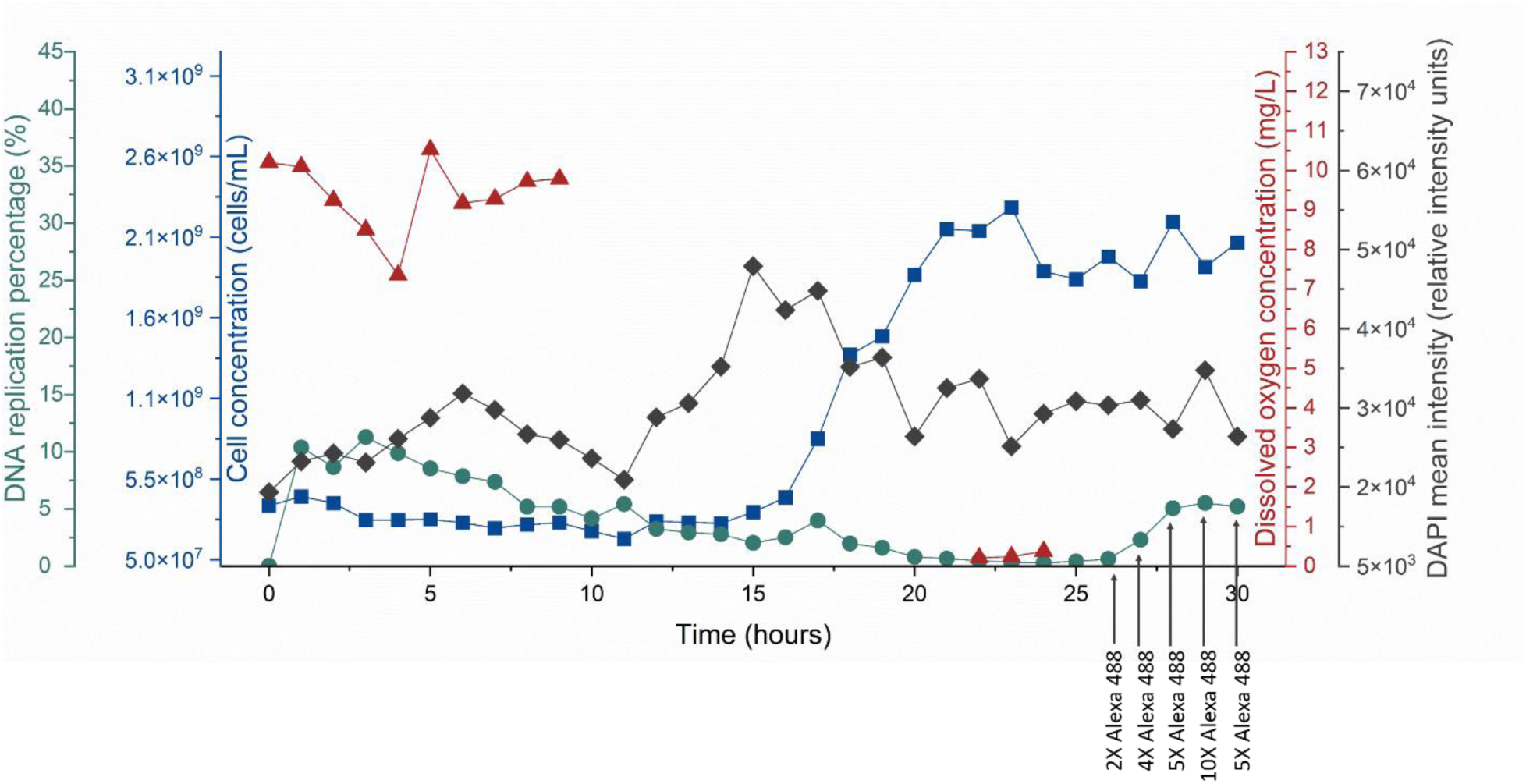
Cell concentration (dark blue), DNA replication percentage (green), dissolved oxygen concentration (red), and DAPI mean fluorescence intensity (dark gray) of *E. coli* cultivated in a 10-mL continuous bioreactor (D= 0.25 h^−1^, T = 30°C, 250 rpm), measured hourly, automatically by the OC-300. Time points from 26 until 30 h indicate the different concentrations of Alexa 488 dye used in the Alexa 488 reaction mix in comparison with the original mix (Table 1).

There are some more findings to discuss using automated on-line flow cytometry. In bioprocesses, understanding the relationship between dilution rate (D) and cell concentration is crucial for optimizing reactor conditions[1,33]. The experiments reveal significant insights regarding the behavior of cell growth under different dilution conditions.

It was observed that the cell concentration during the balanced phase was higher at a lower dilution rate (D = 0.25 h⁻¹) compared to the higher dilution rate (D = 0.5 h⁻¹). The results indicate that the system with D = 0.5 h^−^¹ exceeds the maximum specific growth rates (µ_max_) of the strains used in this study, but this subsequently led to a reduced but stable cell concentration during the balanced phase. Despite dilution rates above µ_max_, no washout of cells was observed, indicating that the cells did not completely exit the reactor.

From a theoretical standpoint, the dynamics of CSTRs (continuous stirred-tank reactor) suggest that washout should occur when the dilution rate surpasses µ_max_, as indicated by a mathematical model of Stephanopoulos and others [34–36]. According to this model, a dilution rate higher than the maximum specific growth rate should result in an immediate washout. However, this was not the case in our observations. In real-world applications, a range of dilution rates beyond µ_max_ exists where washout does not occur, but a reduction in cell concentration is still observed due to suboptimal conditions, which continues until the critical dilution rate (D_c_) is reached. At D_c_, washout occurs, and no further growth is possible[37–40]. This deviation from theoretical predictions can be explained by several parameters that change the environmental conditions in a bioreactor. The higher flow rates can lead to shear stress, nutrient imbalances, changed metabolic balances or other stressors that prevent optimal growth. In addition, the components necessary for cell growth—such as the carbon source, nitrogen, oxygen, and the cells themselves—are never perfectly mixed in a CSTR. These local variations in concentration can affect growth rates even when the overall dilution rate exceeds µ_max_, thereby preventing a complete washout[41–43]. Hence, while the CSTR model predicts washout beyond µ_max_, experimental observations highlight the importance of understanding the complexities of real bioreactor systems. By integrating automated on-line flow cytometry into bioreactor monitoring, comprehensive and real-time insights into cell dynamics and culture conditions can be obtained. Suboptimal growth rates can be detected, allowing proactive intervention to maintain bioreactor performance.

## 5. Conclusion

This study demonstrates that automated on-line flow cytometry is a powerful tool for real-time monitoring and control of bioreactor systems. By providing continuous insights into cell populations, including density, activity, and DNA replication status, this approach overcomes the limitations of traditional offline sampling methods. The integration of a double-staining strategy, using DAPI-FI for total DNA content and Alexa Fluor 488 EdU for active DNA replication, proved effective in capturing cell cycle dynamics, though its applicability remains limited by cost and strain-dependent variability.

The findings emphasize the importance of DAPI-FI as an early and sensitive marker of microbial growth, enabling timely interventions to optimize bioprocess conditions. Additionally, the study highlights discrepancies between theoretical CSTR models and real-world bioreactor behavior, particularly in relation to dilution rate effects and washout dynamics. These deviations underscore the complex interplay of factors influencing microbial growth, including shear stress, nutrient availability, and metabolic imbalances.

Ultimately, the implementation of automated on-line flow cytometry enhances bioprocess efficiency by enabling precise control over culture conditions, reducing variability, and improving product consistency. As bioprocessing industries continue to evolve toward more sophisticated, automated and scalable production methods, this technology represents a valuable advancement for optimizing industrial biotechnological applications.

## Author Contributions

Conceptualization, S.M., K.S.; methodology, J.L.-G, K.S., and M.O.D.S.; investigation, J.L.-G, E.S, H.M., and S.M.; data evaluation, J.L.-G.; writing, review and editing, J.L.-G., S.M., H.H., K.S., and

M.O.D.S. All authors have read and agreed to the published version of the manuscript.

## Acknowledgements

We would like to thank Christine Süring, Johannes Lambrecht and Florian Schattenberg for their help and support during the realization of this work.

