## Supplementary figures for "A double-staining automated flow cytometry method for real-time monitoring of bacteria in continuous bioreactors"

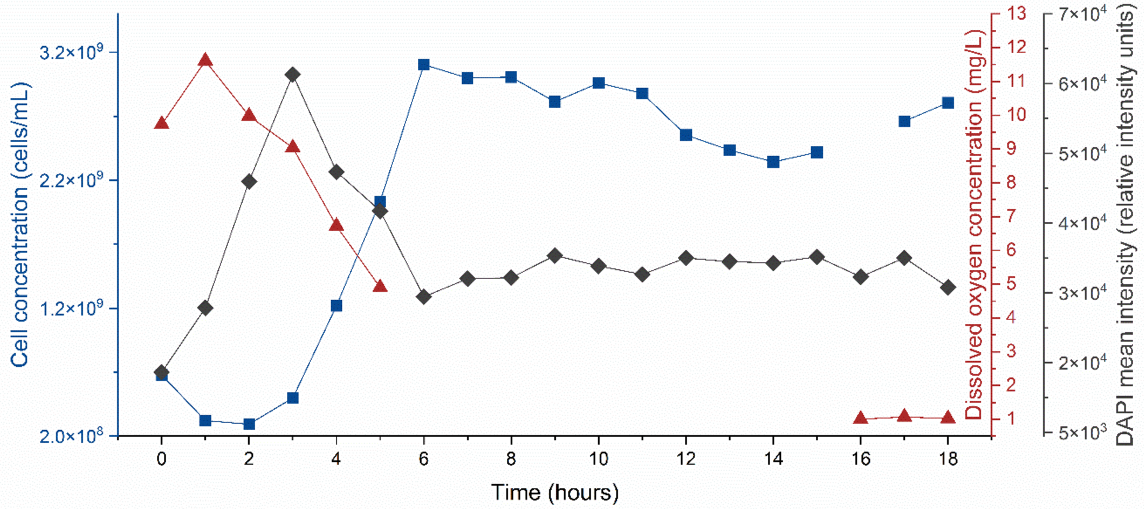


**Figure SI 1**. Automatic on-line cytometric analysis of cell growth in 10 mL bioreactors *E. coli* under continuous balanced conditions at D = 0.31 $h^{-1}$. Cell concentration (dark blue), dissolved oxygen concentration (red), and DAPI mean fluorescence intensity (dark grey) were measured hourly at conditions of D= 0.31 $h^{-1}$, T=30°C, and 250 rpm.

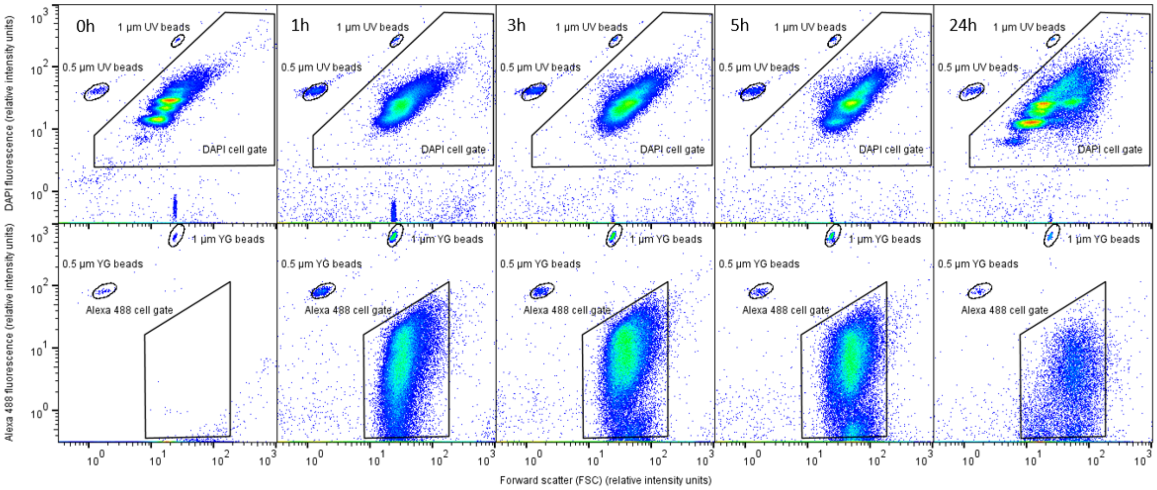


**Figure SI 2.** 2D flow cytometric plots of an *E. coli* culture sampled at various time points. The cells were batch-cultivated in a 24-well plate, manually sampled, and processed. Following fixation using the standard PFA/EtOH protocol[1], the cells were double-stained with DAPI and Alexa 488.

- *Bradyrhizobium* sp. (obtained from Schlechter et al.[2])
- *C. necator* (DSM 13513)
- *E. coli* K12 LE392 (DSM 4230)
- *K. rhizophila* (DSM348)
- *M. rhodesianum* (MB126)
- *P. citronelolis* (obtained from Schlechter et al.[2])
- *P. polymyxa* (DSM36)
- *P. putida* (KT2440)
- *S. melonis* (obtained from Schlechter et al.[2])
- *S. rhizophila* (DSM14405)

**List SI 3.** List of bacterial strains tested for the Alexa 488 and DAPI double staining procedure.
